# Spatial context of cellular heterogeneity in triple-negative breast cancer reveals actionable subtypes

**DOI:** 10.64898/2026.10.10.758236

**Authors:** Zahra Mesrizadeh, Kavitha Mukund, Lixin Yang, Raju Pillai, David Frankhouser, Daniel Schmolze, Susan Neuhausen, Jovanny Zabaleta, Luis Del Valle, Jerneja Tomsic, Yuan Chun Ding, Augusto Ochoa, Lucio Miele, Victoria Seewaldt, Shankar Subramaniam

## Abstract

Triple-negative breast cancer (TNBC) is an aggressive, heterogeneous disease with limited targeted treatment options. Leveraging bulk and spatial transcriptomics and cellular mapping of breast tissue from a large patient cohort, we identified three reproducible TNBC ecosystem subtypes that encode distinct spatial tumor-microenvironment architectures and clinically relevant treatment-response phenotype. Our analyses define: (1) an immune-exhausted subtype associated with immune checkpoint therapy (ICT); (2) a highly fibrotic/hypoxic subtype; and (3) a metabolically reprogrammed subtype characterized by nuclear-receptor and fatty-acid signaling. Spatial profiling further revealed subtype-specific organization of epithelial, immune, and stromal programs across tumor centers and invasive edges. Multiplexed imaging independently supported the contrasting immune-infiltrated and stromal-excluded architectures of the immune-engaged and fibrotic subtypes. The unique markers associated with the subtypes also provide the basis for subtyping and candidate therapeutic vulnerabilities. Our study provides a robust framework for personalized medicine, highlighting that targeting specific spatio-cellular architectures is critical to improving clinical outcomes.

## Introduction

Triple-negative breast cancer (TNBC) is a biologically heterogeneous and aggressive subtype of breast cancer (BC) defined by the lack of estrogen receptor (ERα) and progesterone receptor (PR) expression and no overexpression or gene amplification of human epidermal growth factor receptor 2 (HER2)^1–4^. TNBC occurs with higher frequency in young, parous women, particularly those who self-report as African-American, Black, or African (AA), and in women harboring germline *BRCA1* mutations^5^. Despite major progress in defining TNBC biology, a universally adopted subtyping framework that meaningfully guides clinical decision-making remains elusive. Existing transcriptomic subtypes—such as basal-like (BL), immunomodulatory (IM), mesenchymal (M), and luminal androgen receptor (LAR)—provide informative biological insights^6–10^, but are not typically used in clinical practice, except for androgen receptor (AR) targeting^11–14^. Furthermore, traditional approaches prioritize clustering based on global gene-expression programs, overlooking the cell heterogeneity of the tissue, which dictates the intricate regulatory interactions between malignant epithelial cells and the surrounding tumor microenvironment (TME).

TNBC progression is driven not only by epithelial-intrinsic mechanisms but also by spatially organized stromal and immune niches that dictate selective pressures on tumor evolution^15,16^. The "tumor ecosystem" concept emphasizes that TNBC functions as an interacting, adaptive system composed of malignant epithelial, stromal, and immune cell states that collectively dictate phenotype and therapeutic response^17^. Whereas classical molecular subtyping captures global transcriptomic states, cell-type-informed subtyping is necessary to delineate the underlying tumor ecosystem. Integrating these complementary layers allows for a unified, mechanistically interpretable model of TNBC heterogeneity. While the TME is a central determinant of TNBC progression, prognosis, and therapeutic response, its precise contribution to subtype identity remains incompletely understood. Bulk RNA-seq has historically enabled major advances in TNBC classification, but it averages signals across malignant and non-malignant compartments, masking spatial complexity. Spatial transcriptomics overcomes this limitation by resolving gene expression within anatomically defined tumor and TME regions, providing a mechanistic link between whole-tumor molecular programs and the spatially organized cellular ecosystems that drive TNBC behavior. Recent spatial transcriptomic studies in breast cancer and TNBC have revealed substantial heterogeneity in tumor architecture, including spatially organized immune and stromal niches and their association with tumor behavior^18–22^. However, how patient-level TNBC subtypes defined by cellular composition are reflected in spatial organization of epithelial and microenvironmental programs remains incompletely understood.

Here, we developed a cell-type-driven subtyping framework to identify TNBC ecosystem subtypes from bulk transcriptomics and spatially resolve their epithelial and TME programs across distinct tumor regions. Using 250 pre-treatment TNBC tumors (127 AA, 123 EA), we identified three major ecosystem subtypes: immune-rich with exhausted T-cells, highly fibrotic immune-deficient, and adipocytic subtypes, the last of which features a novel ESR1-mutant subclass. Spatial transcriptomics, multiplexed imaging, and IHC validates the subtyping and provides mechanism-driven phenotypes for each subtype. The spatial organization of cellular states and signaling programs across invasive and non-invasive tumor regions provides insights into the subtype-specific spatial programs and how TME shapes tumor prognosis. These findings offer unique, actionable therapeutic targets and enhance prognostic insights, providing a framework for precision medicine in TNBC.

## Results

### TNBC subtyping based on tissue heterogeneity defined by cell-types and states

We analyzed bulk RNA-seq data from pre-treatment TNBC biopsies of 250 racially diverse women from the Louisiana Tumor Registry (123 EA, 127 AA) (Methods; **Fig. 1a**). Benchmarking with the TNBCtype classifier^8^ confirmed substantial heterogeneity; the classifier excluded 38 tumors from subtype assignment due to modest *ESR1* transcript levels despite IHC-confirmed ER-negativity (<1%), whereas all 38 tumors were retained in our analyses **(Supplementary Fig. 1a)**. Hypothesizing that tissue-level cellular heterogeneity better defines TNBC subtypes, we implemented a deconvolution-based framework utilizing 261 markers across 22 breast-associated cell-types (**Fig. 1b** and **Supplementary Table 1**)^19,23–30^. Unsupervised consensus clustering (CC) of these markers revealed three major metagroups (CC1–CC3), with hierarchical clustering (HC) identifying nine distinct sub-clusters (HC1–HC9; **Fig. 1c**). We focused on the three CC subtypes, as they capture major biological variation, whereas HC groups reflect intra-subtype heterogeneity. CC subtypes were independent of race/ethnicity and age, indicating that the clusters are not driven by patient demographics. While obesity was prevalent across the cohort, only CC3 showed a significant enrichment of Stage 1 tumors **(Supplementary Fig. 1b)**. Using xCell^31^ and EPIC^32^ for cellular deconvolution, we defined CC1 as immune-rich, CC2 as fibroblast-dominant, and CC3 as enriched for adipocytes, fibroblasts, and innate immune cells **(Supplementary Figs. 1c-d)**, prompting deeper analyses into these distinct functional programs.

**Fig 1.**
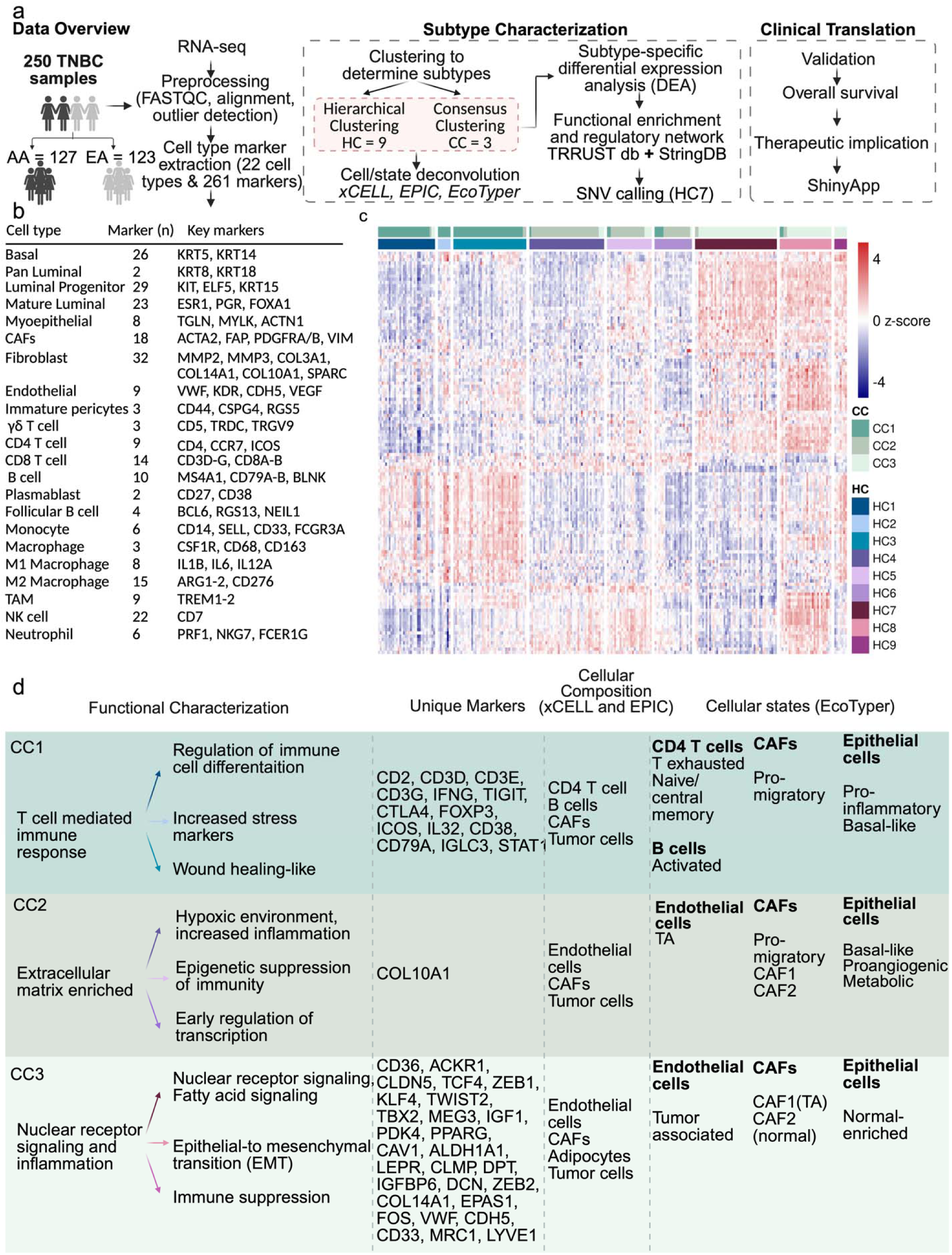
TNBC subtyping and characterization based on cell type marker expression and functional features. **a** Overview of the bulk RNA-seq workflow: from patient to data analysis. Bulk RNA-seq analysis was conducted on fresh-frozen surgical samples from 250 TNBC patients. Bioinformatic analyses were carried out to define cell-type-based TNBC subtypes. **b** The table presents tumor and tumor microenvironment cell types utilized for subtyping LTR TNBCs, along with the number and example of marker genes for each cell type. The complete list of markers is provided in **Table S1**. **c** The heatmap illustrates relative gene expression (Scaled Normalized Values) of cell-type markers within each subtype (ConsensusCluster) and group (HierarchicalCluster) among women with TNBC. This subtyping system categorizes TNBCs into three distinct subtypes and nine groups based on cell-type marker expressions. The visualization highlights the heterogeneity between subtypes and within groups, emphasizing the significance of cell-type markers. **d** This table encapsulates the major functional features associated with TNBC subtypes and groups identified in the LTR TNBC samples, based on their cell-type-associated gene expression. Functional enrichment was performed based on the increased cell-type genes in each subtype and group to unveil underlying mechanisms. (From right to left panel: functional mechanism associated with each subtype, functional mechanism associated with each group, unique cell-type markers in each subtype, combined results from xCELL and EPIC cellular decomposition, and results from EcoTyper cell state analysis are presented). Arrow colors correspond to the group color scheme from the heatmap in **a**.

### Subtype-defining cell-type markers aid functional characterization across TNBC subtypes

To characterize the functional programs of the cell-type-based subtypes, we derived subtype-specific signatures using markers identified through HC (see Methods; **Fig. 1d**). These markers provide structure feature space from which CC1-CC3 differences can be functionally resolved. To further resolve cellular functional states, we applied EcoTyper^33,34^, which identified distinct transcriptional ecosystems across subtypes, linking cell-type and -state diversity to specific activation, exhaustion, and stromal remodeling programs (**Supplementary Fig. 2a**). CC1 exhibited strong immune and stress-response signatures consistent with its immune-rich composition. Key markers highlighted T-cell activation/differentiation, hypoxia/stress regulators, and macrophage-driven ECM remodeling. EcoTyper identified exhausted and naïve/central-memory CD4 T-cell states, activated B cells, pro-migrator cancer-associated fibroblast (CAF) states, and a pro-inflammatory tumor state, together defining an immune-engaged but functionally exhausted microenvironment (**Fig. 1d** and **Supplementary Fig. 2a**). CC2 was enriched for ECM and fibroblast-associated programs, suggesting a stroma/fibrotic dominant phenotype. ECM components, stromal remodeling, innate-immune mediators, and endothelial signatures defined this subtype. EcoTyper identified tumor-associated (TA) endothelial states, pro-migratory CAF, and tumor cells with basal-like and pro-angiogenic states, defining an aggressive, immune-cold, and fibrotic-like microenvironment^35^ (**Fig. 1d** and **Supplementary Fig. 2a**). CC3 was defined by prominent metabolic reprogramming and fatty-acid signaling (FAS), accompanied by WNT regulators, nuclear receptors (NR), and macrophage/ECM-associated genes. EcoTyper identified tumor cells in a normal-enriched state, distinct CAF1/CAF2 states, and TA endothelial cells, supporting a metabolically flexible and stromal-interacting phenotype (**Figs. 1d** and **Supplementary Fig. 2a**). Together, these findings establish three distinct, TNBCtype-independent tumor-microenvironment ecosystems: CC1 (immune-engaged), CC2 (stromal-fibrotic/immune-cold), and CC3 (metabolically reprogrammed). To elucidate the drivers of these phenotypes, we further analyzed the underlying subtype-specific transcriptional networks.

### Subtype-specific transcriptional and regulatory programs in TNBC

To define the molecular programs distinguishing each TNBC subtype, we performed differential expression analysis (DEA) and network analyses. Subtype-specific differentially expressed genes (DEGs) (FDR ≤ 0.05) were identified for each subtype relative to all others and mapped onto curated datasets to construct protein-protein interaction (PPI) and transcription factor-target (TF-target) regulatory networks (**Figs. 2a, 2c-e; Supplementary Fig. 2b)**.

**Fig 2.**
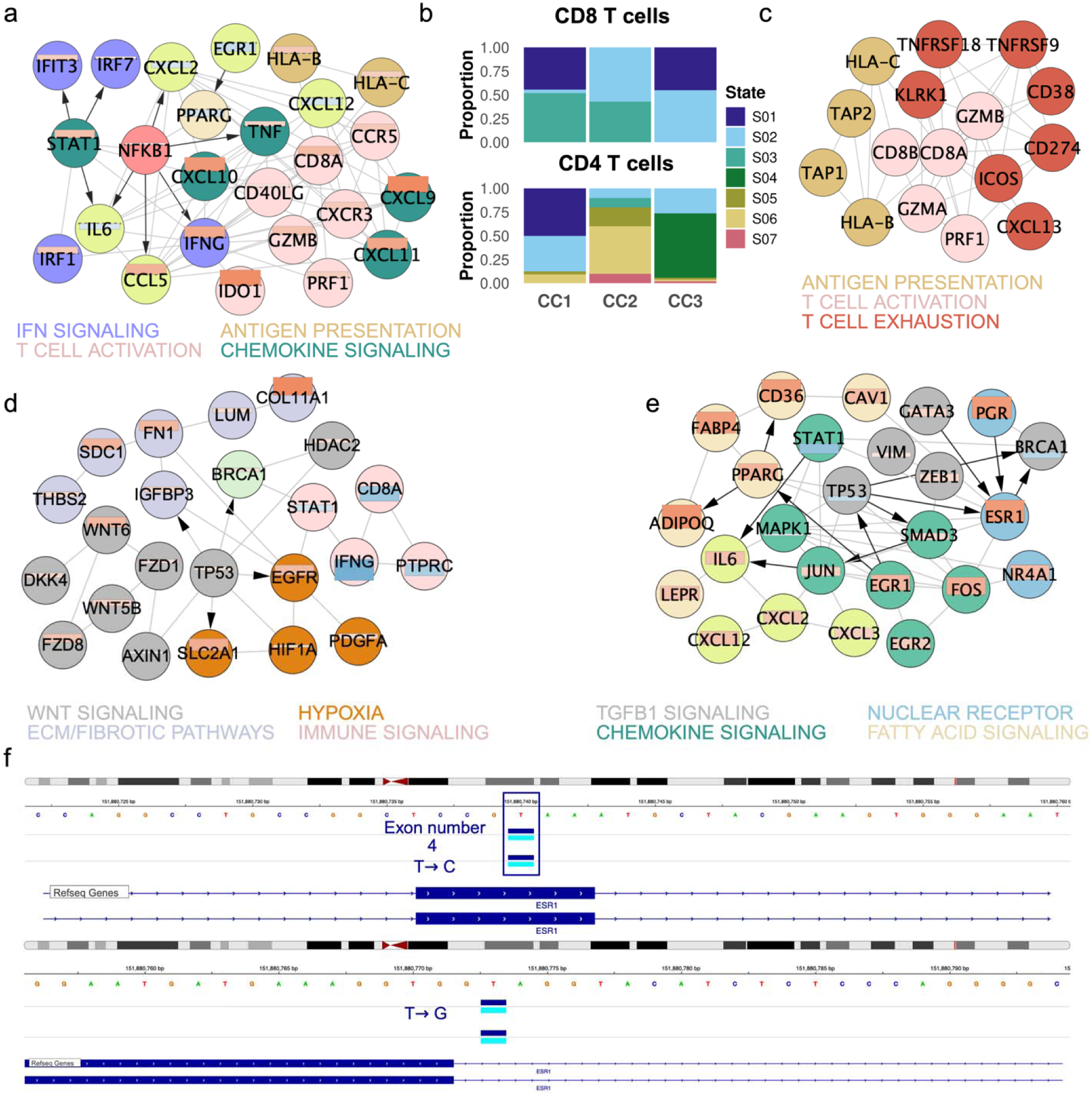
Differentially expressed genes and subtype-specific networks in LTR TNBC. **a** A CC1-specific network highlights immune response pathways. **b** Normalized distribution of T-cell states identified using EcoTyper across subtypes. **c** CC1 specific network highlights exhausted markers and their first neighbors. **d** A CC2-specific network highlights the significance of the extracellular matrix. **e** A CC3-specific network highlights the role of fatty acid, pro-inflammation, and nuclear receptor signaling pathways. The edges present protein-protein interactions, with arrows indicating transcription to target relationships. Nodes of each subnetwork are colored based on the biological mechanism represented in the network, embedded within nodes a heat-bar based on the log2FC value between subtypes: dark blue (log2FC of -2) to white (0) to dark red (2). **f** Representative IGV snapshot showing SNVs on chr6 locations in ESRmRNA^hi^ samples (2 examples for each SNV are shown).

CC1 exhibited strong immune activation governed by *NFKB1*, *STAT1*, *IRF1*, and *IRF7*, accompanied by the suppression of ECM remodeling and fatty acid metabolism. This signature reflects an active *NFKB1*-dominated inflammatory state, integrating antigen presentation, interferon signaling, and proliferative programs, consistent with an immunogenic, danger-sensing microenvironment (**Fig. 2a**). EcoTyper confirmed enrichment of exhausted CD4 /CD8 T cells (**Fig. 2b**). Furthermore, network analysis showed that first neighbors of inhibitory pathways (*ICOS, TIGIT*) were linked to cytotoxicity and antigen processing in CC1 (**Fig. 2c**), defining an immune-engaged but exhausted phenotype potentially responsive to checkpoint blockade. CC2 tumors were defined by ECM-centered remodeling, hypoxia-associated signaling, and non-canonical WNT activation consistent with stromal fibrosis and developmental plasticity. Network analysis identified a *TP53-BRCA1-AXIN1* axis together with WNT5B-associated signaling, supporting coordinated DNA-damage, stromal remodeling, and hypoxic adaptation programs. Furthermore, TP53-induced *IGFBP3* links to a fibronectin-anchored (*FN1*) ECM hub, indicating stromal fibrosis, with EcoTyper defining mesenchymal cells as pro-migratory or tumor-associated CAFs (**Fig. 2d** and **Supplementary Fig. 2a)**. Collectively, these features delineate a transcriptional phenotype of stromal fibrosis and developmental plasticity, consistent with an immunologically cold and aggressive tumor^36^ (**Fig. 2d**). CC3 was characterized by an enriched EMT phenotype driven by *SMAD3*, alongside inflammatory signaling (*IL6*) and distinct metabolic reprogramming (*PPARG*, *CAV1*, *FABP4*), while *MAPK1* was downregulated. Within this cluster, *SMAD3* was associated with a canonical TGFB-EMT axis linked to *ZEB1* and *VIM*, which aligned with AP-1 factors (*JUN*, *FOS*) and *IL6* to establish a TGFB-IL6 inflammatory EMT loop^37,38^. Furthermore, AP-1 linked to *ESR1* upregulation, which connected with *PGR* and *NR4A1*, suggesting aberrant, stress-induced NR activity within a mesenchymal context. Together, these features define CC3 as a transcriptionally plastic, TGFB-inflammatory EMT subtype characterized by metabolic reprogramming and non-canonical steroid receptor signaling (**Fig. 2e**).

### Nuclear receptor programs are enriched in CC3

Given the enrichment of NR-associated transcriptional programs in the CC3, we examined the expression of all 48 human NRs^39^ and identified 23 upregulated receptors, including the entire NR4A family **(Supplementary Fig. 2c)**. NR-centered network analysis revealed a dominant stress-responsive and metabolic hub composed of AP-1 transcription factors and adipogenic regulators, accompanied by suppression of chromatin remodeling, DNA repair, and IFN-STAT signaling modules **(Supplementary Fig. 2d)**. These findings support the presence of a metabolic, NR-associated transcriptional state in CC3 and nominate NR signaling as a potential therapeutic vulnerability for farther investigation.

Although all tumors included in this study were ER-negative by IHC (<1% ER expression), transcriptomic analysis revealed increased *ESR1*, *ESR2*, and *PGR* mRNA expression within CC3, particularly in the HC7 subgroup **(Supplementary Figs. 3a-b)**. Compared with luminal A breast cancers, HC7 tumors displayed distinct transcriptional features and enrichment of arachidonic acid and eicosanoid pathways (**Supplementary Figs. 3c-d**), suggesting that ESR-associated transcription occurs within a broader inflammatory and lipid metabolic context rather than reflecting a canonical luminal phenotype. Together with the TGFB-inflammatory EMT circuitry identified in **Fig. 2e**, these findings characterize CC3 as a transcriptionally plastic subtype associated with metabolic adaptation and non-canonical steroid receptor signaling^40,41^.

To determine whether elevated *ESR1* expression was associated with sequence variation, we examined the ESR1 locus within CC3 tumors and identified two mutually exclusive ESR1-associated variants on chromosome 6, consisting of a synonymous substitution (151880740 T>C) and a splice donor variant (151880773 T>G) (**Fig. 2f**). The synonymous variant was present in 30 of 35 ESRmRNAhi tumors, whereas the remaining five harbored the splice donor variant. Although the functional consequences of these variants remain unclear, their selective enrichment in ESRmRNAhi tumors raises the possibility of an association between *ESR1* sequence variation and aberrant ESR1 transcription that warrants further investigation.

### Subtype-specific spatial architecture of TNBC

To map subtype-specific programs within tissue architecture, we profiled 75 TNBC tumors (20 CC1, 19 CC2, 37 CC3) using GeoMx® DSP across 1,597 segmented regions of interest (ROIs) (**Fig. 3a**). These were then parsed into areas of illumination (AOI) PanCK (epithelial), and PanCK (TME) segments depicting Center (C), invasive edge (E), or distant isolated foci (F) tumor regions chosen by a trained pathologist. Across all subtypes, morphological annotation revealed that regions C and E were primarily composed of invasive epithelia, whereas region F consisted of non-invasive epithelia, including atypia (A) and adjacent normal (J) (**Figs. 3b, c,** and **Supplementary Fig. 4a)**. We identified DEGs between PanCK and PanCK segments, with tumor ROIs showing high cytokeratin enrichment and TME ROIs showing enrichment for immune/ECM genes (FDR < 0.05); **(Supplementary Fig. 4b)**. To characterize the TME, we applied SpatialDecon^42^ to determine cell-type composition (see Methods). Consistent with the high stroma content in TNBC, analysis across all subtypes revealed a dominant fibroblast/mesenchymal population (**Fig. 3d** and **Supplementary Fig. 5a)**. Notably, spatial analysis revealed distinct subtype characteristics. CC1 showed T-cell, B-cell, and myeloid enrichment, particularly at the edge, matching the immune-engaged, interferon/NFKB-driven phenotype from bulk data. Conversely, CC2 and CC3 featured perivascular-like (PVL) populations, with myeloid lineages enriched at foci, indicating distinct stromal and innate immune organization within mesenchymal and fibrotic subtypes (**Fig. 3d** and **Supplementary Fig. 5b)**. Because tumor phenotypes arise from context-dependent functional states within organized ecosystems rather than cell-type identity alone, we used GeoMx to analyze how key signaling pathways—interferon/NFKB, WNT/fibrotic-like, TGFB-IL6 EMT, and NRS metabolic programs—are spatially organized across tumor/TME compartments and across center, edge, and foci regions.

**Fig 3.**
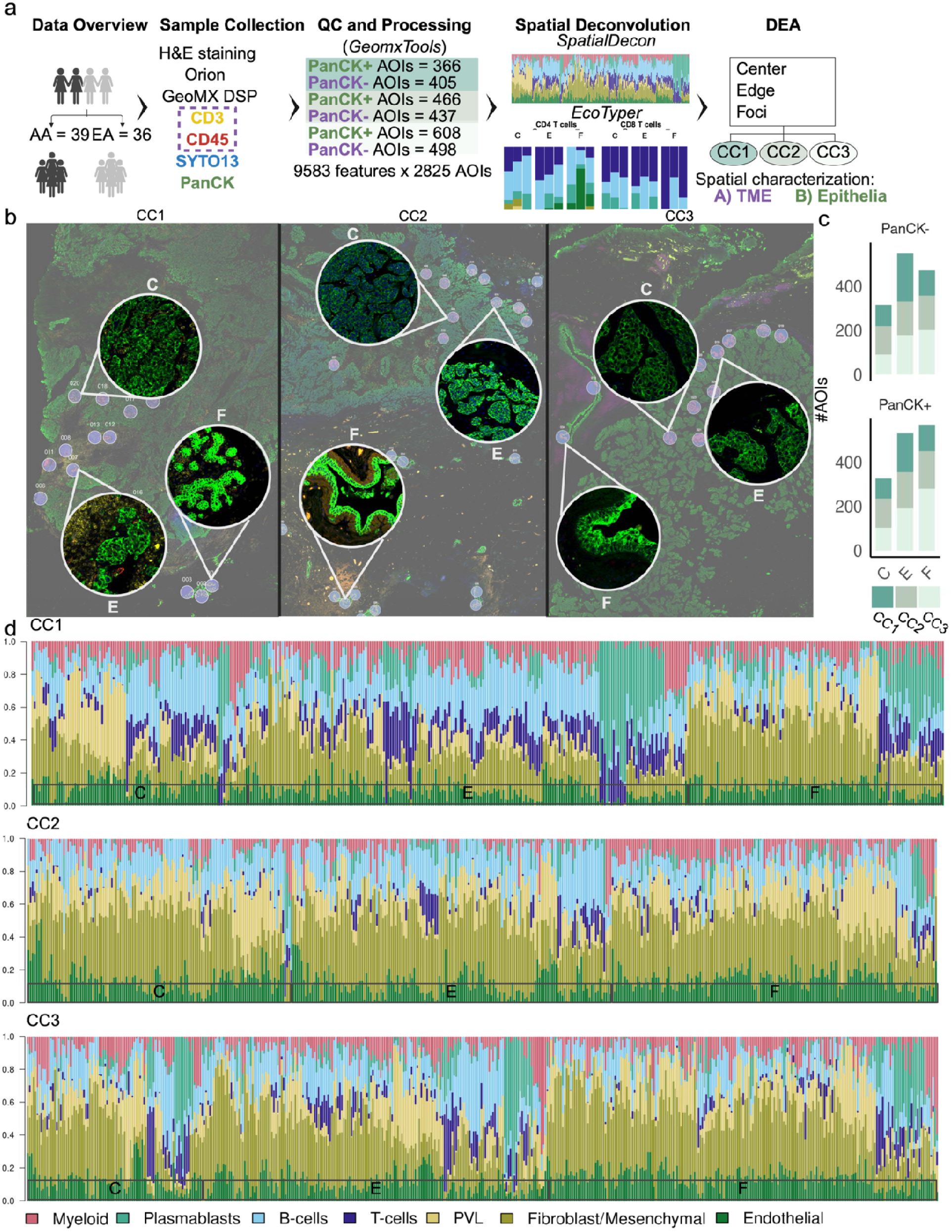
Spatial transcriptomic (ST) characterization of TNBC subtypes. **a** Overview of the ST GeoMx workflow: from patient to data analysis. ST analysis was conducted on fresh-frozen surgical samples from 75 TNBC samples, H&E stained, IF stained for pan-cytokeratin and immune (CD3/CD45), and regions of interest were selected with varying epithelial morphology from different spatial regions. Bioinformatic analyses were carried out to characterize the spatial distribution of cellular composition and signaling programs for epithelial and TME separately. **b** IF imaging examples for each TNBC subtype. ROIs annotated for differing epithelial morphology and spatial regions. ROIs were selected by a pathologist/trained clinician at the tumor interface (E), within the center of the tumor (C), and if available, in regions distant from the center and edge (F, isolated foci). **c** Bar plots showing the distribution of AOIs for each region across TNBC subtypes. **d** The normalized frequency of immune cell types deconvoluted across CC1-CC3 subtypes in PanCK-AOIs generated using SpatialDecon and a custom reference profile matrix. We estimated the abundance of 7 major immune cell types (T-cells, B-cells, plasmablasts, myeloid, PVLs , and fibroblast (mesenchymal cell types) and endothelial cells).

### Spatial organization of cellular states and signaling across invasive and non-invasive regions

To further refine subtype-specific ecosystems, we applied EcoTyper to both the center and edge of invasive ROIs, enabling a systematic characterization of cell-state compositions across tumor and TME compartments. Immune-inflamed tumors (CC1) showed a distinct polarization, with the center dominated by proliferative myeloid and pro-migratory fibroblasts (*SLC29A4* ), while the invasive edge was enriched with exhausted/effector CD4 and CD8 T cells, naïve and activated B cells, cDC1 dendritic cells, classical monocytes, and M1-like macrophages. In contrast, immune-cold tumors (CC2) exhibited a uniform TME rich in CAFs, M2-like foam macrophages, and endothelial cells across both areas. The metabolic subtype (CC3) exhibited a mixed TME with M2 foam-like myeloid cells and mixed fibroblast populations centrally, while the edge featured a heterogeneous, normal-enriched immune-stromal ecosystem (**Fig. 4a**). These TME profiles coincided with distinct epithelial transcriptional states: CC1 (pro-inflammatory), CC2 (basal-like), and CC3 (*AGR2* ) states, consistent with coordinated epithelial-microenvironment organization (**Fig. 4b**). Finally, Orion imaging validated these architectures, confirming high immune infiltration in CC1 and exclusion in CC2. These spatial features were consistently observed across bulk-derived, tumor epithelial, and microenvironmental subtype assignments, supporting the robustness of the molecular classification (**Figs. 4c-f**).

**Fig 4.**
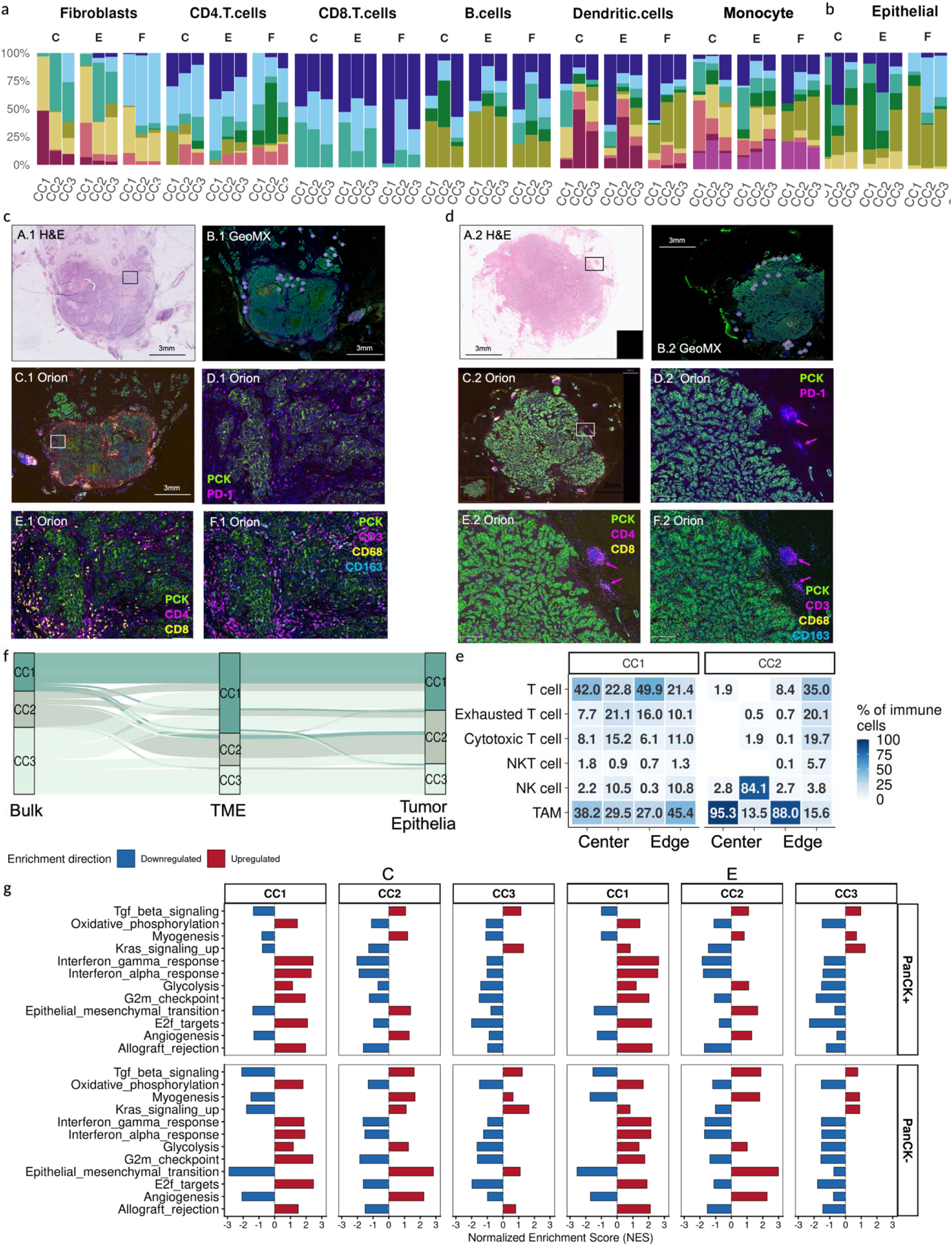
Integrated spatial characterization of tumor and immune ecosystems across TNBC subtypes. **a-b** Normalized distribution of cell states identified using EcoTyper across subtypes and separated by regions (center (C), edge (E), and isolated foci (F)). **c-d** Representative Orion staining of CC1 and CC2 TNBC, respectively: **A** 5 um sections are stained with Hematoxylin and Eosin (H&E). **B** is the corresponding GeoMx image of **A**. **C-F** are the Orion analyses of representative CC1 and CC2 TNBC. In **C**, samples are stained with the Orion panel as outlined in methods and is provided for spatial orientation; the white box illustrates the area detailed in **D**, **E**, and **F**. **D** Depicts co-staining with pan-cytokeratin (PCK) and the immune exhaustion marker, PD-1 (PD-1). **E** illustrates co-staining with PCK and the T-cell markers CD4 (CD4) and CD8 (CD8). **F** depicts co-staining with a) PCK, b) the mature T-cell marker CD3 (CD3), that is critical for antigen recognition and initiation of the immune response, c) CD68 (CD68) a marker expressed on the surface and within macrophages, monocytes, and granulocytes as well as tumor-associated macrophages, and d) the macrophage/monocyte scavenger marker CD163 that can also appear on M2-polarized macrophage (CD163). CC1 image 1 is an immunologically active TNBC with abundant cells that stain positively for CD4, CD8, and CD3; some cells stain positively for PD-1, but many do not. In contrast, CC2 image 2, contains small islands of cells peripheral to the tumor that stain positively for CD4 (**E.2**) and CD3 (**F.2**) but also stain strongly for PD-1 (**D.2**)**. e** Heatmap showing the proportion of immune cell populations (% of total immune cells) across tumor regions (Center and Edge) in CC1 (P1, P2) and CC2 (P3, P4) profiled by Orion multiplex imaging. Cell types were classified based on marker co-expression: T cells (CD3e^+^), Exhausted T cells (CD3e^+^PD-1^+^), Cytotoxic T cells (CD3e^+^GranzymeB^+^), NKT cells (CD3e^+^CD56^+^), NK cells (CD56^+^CD3e^−^), and tumor-associated macrophages (TAM; CD163^+^). Color scale represents the percentage of immune cells per region, ranging from 0% (white) to 100% (dark blue).**f** Contribution of tumor and TME compartments to the TNBC cellular subtypes. The alluvial plot shows the distribution of TNBC subtypes from global bulk RNA-seq, ST tumor and TME pseudobulking**. g** Molecular characterization of invasive tumors (top) and TME (bottom) compartments across TNBC subtypes. The most relevant and significant molecular features from Hallmark are illustrated. FDRs are based on the Wilcoxon rank-sum test. FDR < 0.05 shown; blue = negative association, red = positive association.

To understand how distinct immune-stromal architectures are associated with distinct signaling programs, we conducted gene-set enrichment analysis (GSEA) on both epithelial and TME regions (**Fig. 4g**). Analysis of invasive ROIs revealed distinct pathway activations by subtype (FDR < 0.05). CC1 demonstrated an immune-visible, proliferative phenotype, characterized by high IFNG/IFNa signaling, antigen presentation, and cell-cycle progression in both tumor and TME compartments, with prominent JAK-STAT signaling at the invasive edge consistent with active immune cross-talk. Conversely, CC2 represented an immune-cold, fibrotic-like state, showing downregulation of immune activity and enrichment of EMT, angiogenesis, myogenesis, and TGFB signaling. CC3 was characterized by downregulated cell-cycle programs in both compartments, with specific KRAS pathway upregulation within the TME center (**Fig. 4g**).

Next, we investigated whether regulatory programs observed in invasive regions were also detectable in non-invasive ROIs, including both epithelial (A and J) and paired TME compartments. EcoTyper analysis showed conserved cellular composition across non-invasive TME subtypes, characterized by normal-enriched fibroblasts, endothelial cells, and heterogeneous monocytes, representing a less segregated environment than the invasive front. In the epithelial compartment, cellular states converged on *AGR2* (CC1, CC3) or mixed normal-enriched and *AGR2* (CC2) states, suggesting a shared pre-invasive phenotype (**Fig. 4b**). Despite this structural convergence, GSEA revealed distinct, subtype-specific signaling differences (FDR < 0.05). Non-invasive CC1 showed immune-related enrichment (Notch, IFNa), CC2 exhibited a fibro-inflammatory, immune-suppressed state (TGFB, PI3K), and CC3 showed metabolic stress adaptation (WNT, ROS, UPR) **(Supplementary Figs. 6a-b)**. Together, these findings suggest that functional signaling programs, rather than cellular composition alone, contribute to subtype-specific ecosystem organization and are detectable in non-invasive regions.

### Spatial coupling of regulatory programs within the tumor ecosystem across TNBC subtypes

Spatially resolved analyses revealed that subtype-defining epithelial programs were consistently coupled with complementary immune and stromal states across invasive regions, suggesting coordinated ecosystem-level organization. To define molecular programs associated with invasive regions, we performed subtype-and region-specific DEGs (FDR < 0.05) and PPI-TF-target network analysis (See Methods). While these spatially resolved circuits recapitulated bulk-derived CC programs (**Fig. 2),** they now resolve how epithelial and TME signaling states are spatially coordinated across invasive center-edge regions. Across subtypes, epithelial compartments encoded the dominant regulatory programs that are complementary to TME signaling states. Specifically, the *IFN-STAT1-NFKB* inflammatory axis in CC1, ECM-fibrotic remodeling in CC2, and *MAPK/WNT/TGFB*-driven metabolic plasticity in CC3 were consistently enriched within epithelial-associated programs.

In the immune-inflamed CC1 subtype, tumor cells at the center and invasive edge displayed a high IFN-associated regulatory program, with network analysis identifying an *IRF1-CEBPB-E2F1-DNMT1* axis linking interferon response genes to cell-cycle regulators (**Fig. 5a**). At the invasive edge, an epithelial-intrinsic NFKB/TNF module, involving *STAT1/RELA/TRAF2* activation, was associated with elevated *CCL5* and *CXCL9* expression. This epithelial chemokine output aligned with *CXCR3/CCR1* signaling within the TME, suggesting coordinated immune recruitment and epithelial-immune communication^43^ (**Fig. 5b**). Concurrent upregulation of MHC-I and MHC-II by epithelial compartments further supports an immune-visible epithelial state rather than a passive immune target phenotype. CC2 was characterized by a low-proliferative, stromal-dominant core enriched for collagen-associated genes in TME and reduced proliferative programs within epithelial regions (**Fig. 5c**). In contrast, the invasive edge exhibited compartment-specific signaling rewiring characterized by TME-enriched *FN1-TGFBR1-TGFBI* associated matrix remodeling together with reduced epithelial antigen-presentation and NFKB-associated programs, consistent with a fibro-inflammatory and immune-restricted invasive interace^44^ (**Fig. 5d**). In CC3, tumor epithelial cells exhibited a unique signaling rewiring and metabolic plasticity. At the center, epithelial cells exhibited stress-responsive non-canonical MAPK rewiring, including reduced *MAPK1* expression together with increased *MAP3K1* and *IL1R*^45–47^ (**Fig. 5e**). This was accompanied by increased *RASA1* and *EPHB4*, suggesting vascular-associated signaling^48^. Furthermore, the convergence of WNT (increased *DVL2*, *AXIN2*) and TGFB (induced *TGFBR2* in the TME) was consistent with coordinated epithelial-stromal plasticity, aligned with bulk RNA data (**Fig. 5e**). At the invasive edge, stress-adaptive MAPK signaling persisted through MAP3K1-associated connectivity with *SP1* and *HRAS* despite reduced *MAPK1* expression. Concurrently, an NR-associated metabolic state emerged, defined by epithelial upregulation of *ESRRA* and *NRIP1* and induction of fatty acid metabolism genes (*FABP1*, *CPT1A*). Complementary TME rewiring was also observed, including reduced *RELA* expression alongside increased *AKT3*/*PI3KR3*^49^ and decreased *HSP90AA1*, consistent with reprogrammed PI3K/AKT-HIF signaling states (**Fig. 5f**). Together, these findings define CC3 as a metabolically adaptive and stress-responsive TNBC ecosystem characterized by coordinated epithelial-TME rewiring and non-canonical NR activity despite ER negativity.

**Fig 5.**
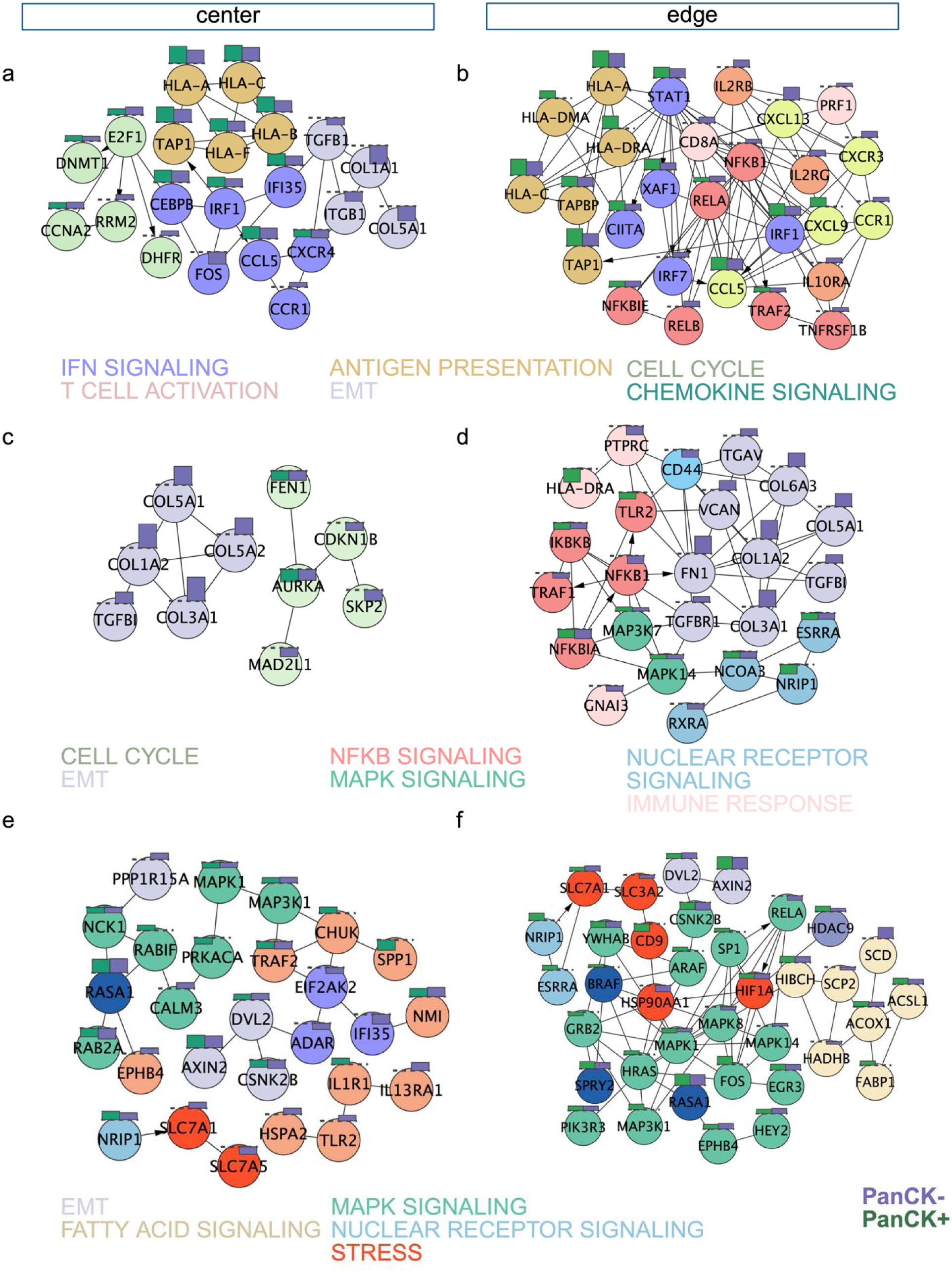
Spatially resolved regulatory networks across TNBC subtypes and invasive regions. **a-b** A CC1-specific network highlights immune response pathways in the center and edge. **c-d** A CC2-specific network highlights the significance of extracellular matrix and EMT in the center and edge. **e-f** A CC3-specific network highlights the role of fatty acid, pro-inflammation, and nuclear receptor signaling pathways. The edges present protein-protein interactions, with arrows indicating transcription to target relationships. Nodes of each subnetwork are colored based on the biological mechanism represented in the network, with bar plots inside nodes colored based on epithelial (green) and TME (purple) and the height based on log2FC.

### Prognostic association and cross-dataset validation of TNBC subtypes

To determine clinical relevance, we analyzed overall survival (OS) in patients with stage I-III TNBC under 63 years old (n_Total_=130, and n_AA_=70 and n_EA_=60). CC1 was linked to better outcomes, while CC2 showed a significantly worse prognosis (*p* < 0.05; **Fig. 6a**). We identified distinct prognostic markers for each subtype: CC1 was defined by *CD244* (favorable) and *SLC4A2* (unfavorable), while CC2 was defined by *GNLY*/*IKZF3* (better survival) and *MIR1262*/*TYR* (poorer survival). CC3 prognosis was driven by genes including *PRF1*/*IL18RAP* (favorable) and *SLFN12L*/*CRTAM* (unfavorable). Some gene associations were context-dependent, requiring further study **(Supplementary Fig. 7a)**.

**Fig 6.**
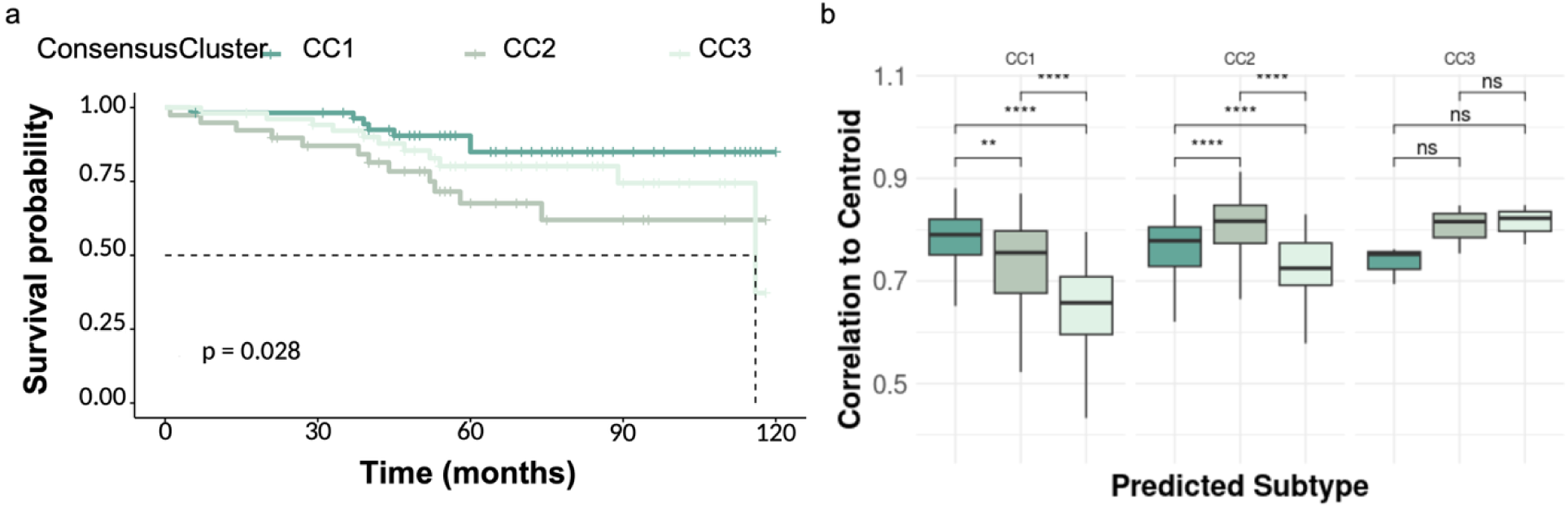
Prognostic relevance and subtype classification robustness of TNBC subtypes. **a** Kaplan-Meier survival analysis stratified by TNBC subtypes (CC1, CC2, CC3) derived from consensus clustering. CC1 tumors show improved overall survival compared to CC2 and CC3, with significant differences across groups (log-rank test, p = 0.028). Tick marks indicate censored observations. **b** Correlation of samples to subtype centroids across predicted subtype assignments. Boxplots show the distribution of correlation coefficients between individual samples and subtype-specific centroids, grouped by predicted subtype (CC1, CC2, CC3). Samples classified as a given subtype exhibit the highest correlation to their corresponding centroid, demonstrating the robustness of subtype assignment. Statistical significance between groups was assessed (Wilcoxon rank-sum test); **** *p* < 0.0001, ** *p* < 0.01, ns = not significant.

The CC1-CC3 framework for TNBC subtyping was validated for robustness and reproducibility using the TCGA TNBC cohort (n = 156) and an independent single-cell TNBC dataset (n = 47)^19,20^, defining three distinct, reproducible ecosystems with unique immune and stromal characteristics. To assign TNBC subtypes to validation data, a centroid-based classification model was performed (**Fig. 6b**), demonstrating reproducibility of the CC1-CC3 framework in an external cohort. Single-cell transcriptomic profiling of 47 treatment-naïve TNBC tumors independently recapitulated the three ecosystems, revealing subtype-concordant cellular programs: CC1 reflected an immune-inflamed but exhausted microenvironment dominated by T and myeloid lineages; CC2 exhibited epithelial-fibroblast proliferative states characteristic of an immune-cold, stromal-rich niche; and CC3 showed hormone-like epithelial plasticity and metabolic remodeling with limited immune engagement **(Supplementary Fig. 7b)**.

To assess clinical relevance, we applied our subtype classifier to the I-SPY2 neoadjuvant trial^50^ data. Although subtype alone did not predict pathological complete response (pCR) across all treatment arms, subtype significantly modified response to the pembrolizumab arm (*p* = 0.027). Specifically, the CC1 subtype showed enhanced benefit from immune checkpoint blockade, while CC2 showed an attenuated response. These findings align with recent studies where pCR cases showed high interferon (IFN) signaling, contrasting with increased pro-angiogenic signaling in pathological non-responders (pNR)^51^. Together, these findings suggest that distinct TNBC ecosystem states exhibit differential therapeutic sensitivities linked to coordinated epithelial-microenvironment programs.

## Discussion

Our findings support a model in which TNBC subtypes represent a conserved spatial ecosystem state arising from coordinated epithelial, immune, and stromal programs rather than tumor-intrinsic transcriptional states alone. By integrating bulk transcriptomics, spatial transcriptomics, multiplexed imaging, cell-state analysis, and regulatory network inference, we identified three reproducible TNBC ecosystems (CC1-CC3) with distinct spatial architectures, signaling programs, and therapeutic associations. Unlike prior TNBC classification systems that primarily define tumor-intrinsic molecular states, our framework resolves subtype identity through coordinated tumor-microenvironment organization across invasive and non-invasive regions.

A central finding of this study is that subtype-defining epithelial programs were consistently spatially coupled with complementary immune and stromal states. Across subtypes, epithelial compartments encoded dominant regulatory programs that aligned with distinct TME signaling architectures, suggesting that TNBC phenotypes are associated with coordinated ecosystem states across tumor and microenvironment. Immune-associated IFN/NFKB signaling in CC1, ECM-fibrotic remodeling in CC2, and stress-adaptive metabolic rewiring in CC3 were all maintained across spatially organized invasive regions, supporting the concept that tumor behavior is shaped by integrated epithelial-microenvironment states rather than isolated cellular compartments.

The CC1 subtype represents an immunologically active yet chronically stimulated ecosystem, in which spatial epithelial-immune interactions are associated with simultaneous immune visibility and exhaustion. Transcriptomics revealed enriched IRF1/NFKB activity and identified *CD244* as a complex, dual-role prognostic marker associated with both cytolytic activity^52^ and exhaustion^53–55^. Spatial analyses demonstrated an "immune-hot" center, characterized by heightened MHC-I processing, IRF1-driven signaling, and elevated cell cycle activity. Meanwhile, elevated *DNMT1* raises the possibility that epigenetic regulation contributes to the immune state observed in CC1, indicating a potential role for epigenetic therapeutic strategies aimed at restoring immunogenicity^56–58^. At the invasive edge, CC1 presents a complex, heterogeneous immune-stromal network. Spatial analyses further suggested a compartment-specific role for NFKB signaling, with immune-associated activation within the TME and epithelial-associated IFNG-responsive chemokine programs^59–61^. Notably, the *CXCL9-CXCR3* axis was prominently enriched, supporting its potential relevance as both a prognostic biomarker and therapeutic intervention in breast cancer^43,62–65^. The differential association between subtype and pembrolizumab benefit observed in the I-SPY2 clinical trial further supports the potential clinical relevance of the inflammatory yet epigenetically constrained CC1 ecosystem and its potential susceptibility to immune checkpoint blockade.

Our study identifies CC2 as an immunologically cold, fibroblast-dominated TNBC ecosystem defined by dense ECM organization, hypoxia-associated signaling, and stromal plasticity, in contrast to the immune-engaged CC1 subtype. Instead of supporting proliferative epithelial growth, this subtype prioritizes structural reinforcement, associated with an aggressive, treatment-resistant TNBC phenotype^66–70^. Spatial analyses demonstrated a collagen-dense tumor core together with a more dynamic invasive edge. These findings align with previous studies suggesting that ECM-dense regions may restrict immune infiltration^66^, while the invasive edges retain stromal plasticity. Early TGFB activation was associated with an aggressive, pro-fibrotic environment, limiting immune infiltration, and may contribute to the attenuated pembrolizumab benefit observed in the I-SPY2 trial. These findings suggest that ECM remodeling and TGFB-associated signaling represent candidate therapeutic vulnerabilities for overcoming immune resistance within this subtype.

CC3 represents a distinct immune-excluded TNBC ecosystem in which both tumor and TME compartments preferentially exhibit stress-adaptive and metabolically flexible programs over proliferative signaling. At the molecular level, this subtype showed reduced canonical ERK-associated signaling together with enrichment of stress-responsive JNK/p38-associated programs, consistent with adaptation to inflammatory and metabolic pressure^71,72^. Bulk RNA-seq identified a distinct NRS marked by *ESR1/PGR-associated* DEGs, suggesting a luminal epithelial plasticity despite ER negativity^73^. Spatial profiling further clarified that this apparent luminal-like signature was associated with metabolic adaptation rather than canonical hormone signaling. Enrichment of *ESRRA*, *NRIP1*, and fatty acid metabolism programs suggests a coordinated metabolic state characterized by altered mitochondrial and lipid programs. At the invasive edge, these metabolic pathways converged with WNT and TGFB-associated signaling and were associated with epithelial plasticity and stromal remodeling. Together, these findings support CC3 as a metabolically adaptive and stress-responsive ecosystem distinct from the fibrotic architecture of CC2 and nominate NR, metabolic, and WNT/TGFB-associated pathways as candidate therapeutic vulnerabilities.

Although TNBC studies frequently focus on invasive regions, analyses of non-invasive compartments provided complementary insights into subtype-associated ecosystem organization beyond the invasive tumor. We found that non-invasive regions retained subtype-specific signaling programs despite greater convergence in cellular composition. Notably, CC1 regions demonstrated enrichment of IFN-associated signaling and CXCL13 expression, consistent with an immune-associated state. Conversely, CC2 exhibited TME-restricted TGFB-associated signaling, suggestive of stromal activation, while CC3 displayed metabolic programs involving WNT and adipogenesis signatures. Collectively, these findings suggest that molecular features associated with the three TNBC ecosystems are detectable beyond histologically invasive regions and suggest that subtype-associated ecosystem organization extends across distinct tissue compartments.

Our study establishes a framework for understanding TNBC heterogeneity through patient-level ecosystem subtypes that integrate tumor-intrinsic and microenvironmental programs. By linking these subtypes to their spatial organization across tumor compartments, our findings extend bulk transcriptomic classification to reveal distinct epithelial-TME architectures and candidate therapeutic vulnerabilities. Together, these results provide a foundation for spatially informed patient stratification and therapeutic investigation in TNBC.

## Methods

### Patient cohort and RNA-seq processing

As described in Hossain et al.^73^, 3188 women with TNBC from the Louisiana Tumor Registry (LTR) were selected, of which, 253 TNBC pre-treatment samples were subject to RNA-sequencing. After quality control, 250 samples (127 African American, 123 European American) were retained for downstream analyses. These 250 samples are henceforth referred to as LTR TNBC. Genetic ancestry was available for 227 samples based on 163 ancestry-informative SNPs, as previously described^74^. The quality of raw sequencing data was evaluated using *fastqc v0.11.9* and summarized for downstream analysis using *multiQC*. RNA read mapping and alignment were performed on the human reference genome GRCh38.p13_2021v using *Rsubread v2.10.5*, to produce a count matrix containing 44,715 genes and 250 samples, which were filtered for low-count genes. The resulting data matrix containing 23,264 genes across 250 samples was used for all downstream analysis. Count matrix normalization and differential analysis were performed using *DEseq2 v1.36.0* in R, as outlined in their vignette^75^. Differentially expressed genes (DEGs) for every comparison were called at a Benjamini-Hochberg false adjusted *p* < 0.05. Z-scores for each gene were computed, and genes with a standard deviation (SD) less than ± 2 were filtered to eliminate genes with low biological significance. A total of 18,308 genes passed our significance threshold, and their normalized expression was used as input for visualizations. Heatmaps were generated using the *pheatmap v1.0.12* library in R. The colors on the heatmap represent the normalized expression count scaled row-wise using the built-in scale function available through *pheatmap*.

### Collective Cell Type Markers and subtype Discovery

Cell type markers from twenty-two cell types expected to play a role in breast cancer tumors and tumor microenvironments were identified in the literature **(Supplementary Table 1)**. Group-specific clusters were detected using a hierarchically clustered tree with the dynamic tree cut algorithm at a cut height of 1.3. *ConsensusClusterPlus v1.76.0*^76^ was additionally employed to identify metagroups in an unsupervised manner, with 1000 iterations and the optimal number of clusters (subtypes) (*k*) at 3, determined using the cumulative distribution function (CDF) curve. To identify gene markers uniquely defining each subtype, we converted the z-scores to percentile values using the *pnorm* function in *stats v4.2.0* library in R, adjusting for an exact mean and SD of each sample. Unique subtype gene markers were defined as genes expressed in the top 20% in at least 50% of the subtype’s population. *VennDiagram v1.7.3* and *ggvenn v.0.1.10* were used for all overlap visualizations.

### Spatial transcriptomic profiling of TNBC using the GeoMx Digital Spatial Profiler

Spatial profiling was performed on 75 pre-treatment FFPE tumors using GeoMx-DSP^TM^ (DSP; NanoString Technologies). Tissue sections (5 μm) were stained with H&E and hybridized with the GeoMx Whole Transcriptome Atlas (HuWTA) probe set (∼18,000 genes). Slides were stained with GeoMx Solid Tumor TME and Nuclear Morphology kits, and circular ROIs (300 μm; ≥3 per region) were selected at tumor center, tumor edge, and non-invasive foci. Automated segmentation defined PanCK epithelial and PanCK stromal compartments. UV-mediated oligo release was guided by digital micromirror device (DMD) masks, and oligos were collected per AOI for library preparation using GeoMx SeqCode primers. Libraries were pooled, purified (AMPure XP), assessed by Bioanalyzer, and sequenced on Illumina NextSeq2000 (2×27 bp, ∼100 reads/μm²). FASTQs were processed into DCC files using GeoMxNGSPipeline.

Quality control and normalization were performed using *GeomxTools v3.0.1*. Samples were filtered based on recommended criteria (min nuclei = 40, min area = 1,000 μm², LOQ = 2), retaining genes detected above background in ≥5% of AOIs. AOIs with <5% detectable genes were removed. Q3 normalization as recommended by GeoMx after gene filtering and transformed into log values for downstream analysis, when necessary (log_q). The comparison of interest dictated whether p-values or adjusted p-values (FDR) were chosen for thresholding and identifying differentially expressed genes. Differential analysis was performed using a linear mixed model. Comparisons of interest once again dictated whether an LMM was performed with a random slope (across slides) or with a random slope and intercept (across slides with nested random intercepts).

### Differential analysis using Linear Mixed Models

Differential expression analysis was performed using linear mixed-effects models (LMMs) implemented through the *GeoMxTools* framework, following the recommendations outlined in the *GeoMxTools* vignette. LMMs are well-suited for spatial transcriptomic data because they explicitly account for the hierarchical structure of GeoMx profiling, where multiple regions of interest (ROIs) obtained from the same slide or patient are not independent observations. For this study, we employed an across-slide LMM design with nested random intercepts, appropriate for comparisons spanning multiple slides and patients. This approach models biological and technical structure by incorporating patient-level and slide-level random effects while testing fixed effects associated with the biological comparison of interest (subtypes CC1, CC2, CC3, and regions center, edge, isolated foci). This model structure allowed us to adjust for both repeated sampling per tissue block and variability introduced by slide processing. Analyses were performed separately for PanCK epithelial and PanCK TME segments and further stratified by spatial region. For invasive analyses, ROIs were subset into center (C) and edge (E), whereas non-invasive analyses used isolated foci (F), subset into atypia (A) and adjacent normal (J) epithelial compartments and their matched TMEs.

### Orion Analysis

Consecutive tissue sections were cut from eight (2 CC1, 3 CC2, 3 CC3) of the same tissue blocks profiled by GeoMx and were processed using the Orion multiplex immunofluorescence protocol (Akoya Biosciences) with a custom 20-marker panel. Orion images were processed and analyzed using QuPath v0.5.1. Regions were manually annotated on each image using square annotations matched to the diameter of the GeoMx ROIs (300 µm), allowing corresponding GeoMx ROIs to be manually registered and classified as Center, Edge, or Isolated Foci regions. Cell segmentation was performed using the QuPath cell detection workflow with default settings, and intensity features were calculated for each biomarker stain channel using the add intensity features function. For each channel, cell-level mean intensities were modeled using a two-component Gaussian mixture model. The intersection point between the means of the two fitted distributions was used as the positivity threshold, and cells with mean intensity values above this threshold were labeled positive for the corresponding biomarker. Marker positivity calls were then combined to assign cell phenotypes, with T cells defined as CD3e^+^, Exhausted T cells as CD3e^+^PD-1^+^, Cytotoxic T cells as CD3e^+^GranzymeB^+^, NKT cells as CD3e^+^CD56^+^, NK cells as CD56^+^CD3e^−^, and tumor-associated macrophages (TAM) as CD163^+^. The number of cells assigned to each phenotype was quantified within each individual region and compared between regions using the Wilcoxon test.

### Enrichment, visualization, and reference network generation

Gene set enrichment analysis (GSEA) was performed using the MSigDB Hallmark collection (category H) obtained via the *msigdbr* (R package v25.1.1). Differentially expressed genes for each contrast (CC1, CC2, CC3) were ranked using a composite score defined as the sign of the log2 fold change multiplied by the -log10 adjusted P value. Ranked gene lists were analyzed using the *fgsea* algorithm (R package v1.22.0) with minimum and maximum gene-set sizes of 15 and 500, respectively. For each subtype, normalized enrichment scores (NES) and adjusted P values were computed, and significant pathways were defined as those with FDR < 0.05. For optimized visualization the terms have been shortened, and selected pathways were shown to avoid repetition. Additional GO and functional enrichments were performed using *clusterProfiler*^77^ or *Enrichr*^78^ in R. A high confidence (>0.7) STRING database v12.0^79^ human protein-protein interactions (PPI) containing 15,588 proteins and 236,930 interactions. Additionally, TF-target data downloaded from TRRUST-db^80^ was mapped onto the PPI interaction network to generate a custom PPI/TF network. Bulk-seq derived subtype-specific PPI/TF-target networks were extracted using subtype-specific DEGs, resulting in the CC1-specific network containing 470 proteins with 1,140 interactions, the CC2-specific network containing 276 proteins and 557 interactions, and the CC3-specific network containing 859 proteins and 4,564 interactions. GeoMx derived subtype-and region-specific PPI/TF-target networks resulted in CC1 Center 237 proteins and 534 interactions, CC1 Edge 466 proteins and 1495 interactions, CC2 Center 53 proteins and 22 interactions, CC2 Edge 328 proteins and 508 interactions, CC3 Center 230 proteins and 236 interactions, CC3 Edge 520 proteins and 1299 interactions. These networks were further customized to capture the functional mechanisms underlying each subtype by selecting all DEGs, then reducing to the TF-targets and GSEA enrichment associations. All networks were constructed, annotated, and analyzed in Cytoscape ^45^.

### Spatial cell-type deconvolution using SpatialDecon and a TNBC-specific reference matrix

To improve cell-type deconvolution of PanCK segments, a TNBC-specific reference matrix was constructed using scRNA-seq data from Kumar *et al*.^82^ 100,064 cells were downloaded, and 30,719 non-epithelial TNBC cells were retained. Seven major cell types (B cells, T cells, Myeloid, Endothelial, PVL, Plasmablasts, Fibroblast/Mesenchymal) were used to build a reference profile using *SpatialDecon v1.1* (minCellNum=10, scalingFactor=1, minGenes=100). Deconvolution was performed using *RunSpatialDecon*, yielding cell abundance estimates per AOI.

### EcoTyper

TPM-normalized GeoMx expression (before Q3 normalization) for each segment was input to EcoTyper to recover cell states using the ecotyper_recovery_bulk.R pipeline. Cell-state abundances were multiplied by SpatialDecon-estimated cell-type fractions to produce state-weighted abundance scores. State annotations followed the original EcoTyper publication^33^.

### Survival Analysis

Overall survival (OS) was compared using Kaplan Meier plots and log-rank tests using *survival v3.7-0* and *survminer v0.4.9* in R. OS was defined as the time from diagnosis to death from any cause.

### Variant Calling

Single nucleotide variants (SNVs) and indels were called in all samples from CC1, CC2, and CC3 subtypes, focusing on the ESR1 chromosome locus using *FreeBayes* v1.3.5. Variants were filtered based on quality (QUAL>20) and depth (INFO/DP>100). The filtered variants were annotated using snpXplorer to identify their functional consequences.

### RShiny App

The subtyping approach developed in this study is deployed as an interactive RShiny app for prospective samples and is publicly available via [https://www.tnbcworkbench.org/shiny/TNBCsubtyping].

### Validation Strategy

TCGA-BRCA raw count data for 846 tumors were downloaded using TCGAbiolinks in December 2022 and used for validation of our patient subtyping approach, using the unique markers identified. Samples were characterized into Luminal A (n = 586), Luminal B (n = 107), and TNBC (n = 153) based on their IHC/FISH receptor status. Single-cell TNBC data were accumulated from publicly available studies (Xu *et al*.^19^, Bassez *et al*.^20^) with total of 47 pre-treatment TNBC patients. Normalized gene expression data for testing datasets were correlated (Pearson) to the centroids of the unique gene markers for each TNBC subtype identified above. Each testing sample was assigned to the TNBC subtype with the highest correlation.

## Acknowledgments

The authors wish to thank Professor Jason Swedlow and Dr. Josh Titlow from the Wellcome Leap Foundation for invaluable discussions. The research reported here was supported by a grant from the Wellcome Leap Delta Tissue Program to S.S. and V.S. This work was also partially supported by grants from the National Institutes of Health to S.S. (OT2-OD030544, OT2-OD036435, and R01-CA282657) and the National Cancer Institute/National Institutes of Health to L.M. and A.O. (P20 CA 233374).

## Author Contributions

Z.M. and S.S. designed the systems biology analyses, with Z.M. performing the majority of analyses presented within the paper with guidance from K.M. D.F. performed all the necessary processing and analysis of the image files (Orion and GeoMx). L.Y., J.T., and R.P. conducted the GeoMx assays and generated the data. D.S. and V.S. were lead pathologists in GeoMx study, with V.S. identifying and annotating all ROIs considered within this study. J.Z., L.M., and A.O. were involved in collecting the patient cohort samples used in this study. S.N. and Y.C.D. was responsible for genetic ancestry information. L.D.V. was the lead pathologist at LSU. V.S. was responsible for all patient sample choice/design for the study. S.S. designed and supervised the entire study. Z.M. wrote the first draft of the manuscript, with revisions by K.M. and S.S., as necessary. All authors have read and agreed to the submitted version of the manuscript.

## Data and code availability

The datasets are available through GEO under SuperSeries accession GSEXXX. The code is available on GitHub (https://github.com/zahramesri/TNBC_Subtyping).

